# Deep conservation of *Hid*-like RHG gene family homologs in winged insects

**DOI:** 10.1101/2021.03.30.437773

**Authors:** Markus Friedrich

## Abstract

Together with *sickle* (*skl*), the *Drosophila* paralogs *reaper* (*rpr*), *head involution defective* (*hid*), and *grim* (RHG) control a critical switch in the induction of programmed cell death. RHG homologs have been identified in other dipteran and lepidopteran species but not beyond. Revisiting this issue with a “taxon hopping” BLAST search strategy in current genome and transcriptome resources, I detected high confidence RHG homologs in Coleoptera (beetles), Hymenoptera (bees+wasps), Hemiptera (true bugs), termites, and cockroaches. Analyses of gene structure and protein sequence conservation revealed a shared ancestral splicing pattern and highly conserved amino acid residues at both the N- and C-terminal ends that identify *hid* as the most ancestrally organized RHG gene family member in *Drosophila*. hid-like RHG homologs were also detected in mosquitoes, redefining their *michelob_x* (*mx*) genes as an expansion of derived RHG homologs. Only singleton homologs were detected in the large majority of other insect clades. Lepidopteran RHG homologs, however, stand out by producing an evolutionarily derived splice isoform, identified in previous work, in addition to the newly detected *hid*-like isoform. Exceptional sequence diversification of select RHG homologs at the family- and genus-level explain their elusiveness in important insect genome model species like the red flour beetle *Tribolium castaneum* and the pea aphid *Acyrthosiphon pisum*. Combined, these findings expand the minimal age of the RHG gene family by about 100 million years and open new avenues for molecular cell death studies in insects.

## INTRODUCTION

Programmed cell death results from the unleashed activity of caspases, a deeply conserved gene family of cysteinyl aspartate proteinases. First characterized for their executive role in programmed cell death in the nematode *Caenorhabditis elegans* (Yuan et al. 1993), subsequent studies in other model organisms, i.e. *Drosophila* and mice, uncovered the functional conservation of caspases as executive forces in the programmed cell death pathway (Crawford et al. 2012). However, the mechanisms in control of appropriate caspase activation have been found to utilize both conserved and diverged mechanisms. In mammals, for instance, mitochondrial signals and members of the Bcl2 gene family are in control of caspase activation (Cory and Adams 2002). In *C. elegans*, a similar, but less complex regulatory protein machinery appears to be in place (Lettre and Hengartner 2006). In Drosophila, caspases are constitutively expressed but blocked by default through the physical interventions by members of the Inhibitor of Apoptosis (IAP) gene family (Wang et al. 1999). Pending developmental cues or cellular stress conditions, this block is relieved by the small protein products of the RHG gene family (Conradt 2009), which includes the name-giving paralogs *reaper* (*rpr*), *head involution defective* (*hid*), *grim*, besides *sickle* (*skl*) (White et al. 1994; Grether et al. 1995).

*Rpr* was the first characterized Drosophila RHG gene family member (White et al. 1994), followed by *hid* (Grether et al. 1995), *grim* (Chen et al. 1996), and *skl* (Christich et al. 2002; Srinivasula et al. 2002; Wing et al. 2002). Subsequent efforts of identifying homologs in newly available Drosophila species genome drafts revealed the conservation of all four clustered genes in drosophilid Diptera (Zhou 2005). Similar efforts to find RHG homologs in the first genome draft of the Malaria vector mosquito species *Anopheles gambiae*, however, were unsuccessful (Zhou 2005). At the same time, the comparative analysis of Drosophila RHG homologs corroborated the high conservation of the N-terminal IBM (IAP-binding motif) sequence (Shi 2002; Berthelet and Dubrez 2013): A-[KTVI]-[PAE]-[FEISY]. This finding was consistent with the subsequent discovery that the inhibitory binding of RHG homologs to IAP proteins was dependent on the N-terminal amino acid residues (Zachariou et al. 2003). Comparative sequence analyses further suggested the existence of IBM subtypes (Zhou 2005) and the presence of a second, putatively shared motif called Trp-block or Grim Helix 3 (Wing et al. 2001; Clavería et al. 2002). This progress notwithstanding, the challenge of identifying RHG genes on the basis of very limited sequence conservation culminated in the cautionary statement that even the relatedness of the Drosophila paralogs was only tentatively supported by sequence similarity (Zhou 2005).

Today, caspase and IAP genes have been identified in a wide range of insect species (Ribeiro Lopes et al. 2019), but the search for RHG homologs has thus far been only successful in dipteran and lepidopteran species (Yoo et al. 2017). In Diptera, RHG homologs have been analyzed in the blowfly species *Lucilia cuprina* and *Lucilia sericata* (Chen et al. 2004; Edman et al. 2015), the Carribean fruit fly species *Anastrepha suspensa* (Tephritidae) (Schetelig et al. 2011), and, most recently, the scuttle fly *Megaselia scalaris* (Yoo et al. 2017). In the Lepidoptera, RHG homologs have been studied in the silkmoth *Bombyx mori* and the fall armyworm *Spodoptera frugiperda* (Bryant et al. 2009; Wu et al. 2013; Shu et al. 2020). The existence of RHG homologs outside the Lepidoptera and Diptera, however, has remained elusive. It is possible that the RHG gene family originated in the lineage to the last common ancestor of the Lepidoptera and Diptera, which are relatively closely related insect orders (Misof et al. 2014). The short sequence lengths and low sequence conservation of RHG genes, however, are suspected to limit the detectability of distantly related homologs (Zhou 2005; Ribeiro Lopes et al. 2019). This issue may be exacerbated by the continued scarcity of cell death pathway studies in other insect models (Colella et al. 2018; Ribeiro Lopes et al. 2019; Lopes et al. 2020). Confirming these concerns, I here report the results from searching current genome and transcriptome databases a taxon hopping search strategy, which recovered RHG homologs from a substantially wider range of winger insect orders by a taxon hopping search strategy.

## RESULTS

### RHG homologs from an extended range of winged insects

Initial searches for RHG homologs outside Diptera and Lepidoptera were conducted using the silkworm RHG homolog *IAP-binding motif 1* (*IBM1*) (NP_001159813.1) as query in BLASTp searches against the NCBI nr database with Diptera and Lepidoptera excluded from the taxonomic search range (Bryant et al. 2009). This effort yielded low confidence hits against candidate homologs in the hemipteran species *Bemisia tabaci* (LOC109029550; e-value = 0.021), *Nilaparvata lugens* (LOC111048366; e-value = 0.005), and *Laodelphax striatellus* (RZF36208.1; 0.005). All of these sequences started with the RHG homology-defining IAP binding motif (IBM) (Zhou 2005), were less than 300 amino acids long, and returned IBM1 as single best hit when reBLASTed against the silkworm protein sequence database.

As the presence of RHG homologs in hemipteran species predicted the conservation of the RHG family throughout the Holometabola, I used the newly identified hemipteran sequences as queries in clade-specific BLAST searches for homologs in the Coleoptera (beetles) and Hymenoptera (bees+wasps). This approach produced significant hits in over 50 hymenopteran species, 21 of which were compiled for detailed analysis (Supplementary data file 1) and five coleopteran species. Among the latter, a notable absence was the flour beetle *Tribolium castaneum*, which represents the first coleopteran genome draft that has since been improved by a number of revisions (Richards et al. 2008; Herndon et al. 2020). I therefore continued to BLAST search for additional coleopteran RHG homologs using the newly detected coleopteran homologs as seed queries. One of them, i.e. the putative RHG homolog of the Emerald ash borer *Agrilus planipennis* (XP_018330969.1), detected the protein product of *T. castaneum* locus LOC103313285 (XP_008194456.1) as a candidate RHG homolog with an e-value of 0.05. ReBLAST of the *T. castaneum* LOC103313285 protein sequence against the conceptual *A. planipennis* proteome returned the putative *A. planipennis* RHG homolog as the best matching hit. Using the putative *T. castaneum* RHG homolog as a query against coleopteran transcript and protein sequence databases expanded the compilation of coleopteran RHG sequences to 25 (Supplementary data file 1). This included two further darkling beetle family homologs (*Asbolus verrucosus* and *Zophobas atratus*) and five additional homologs from families in the Tenebrionoidea (Supplementary data file 1).

Similar “taxon-hopping” BLAST searches unearthed high confidence RHG homologs in a total of 19 hemipteran species including aphids as well as in three representatives of the Dictyoptera: The German cockroach *Blattella germanica* (PSN40724) and the termite species *Cryptotermes secundus* and *Zootermopsis nevadensis* (Fig. 1). Extensive searches outside these taxa in pancrustacean and other invertebrate databases did not return candidate RHG homologs.

**Figure 1:**
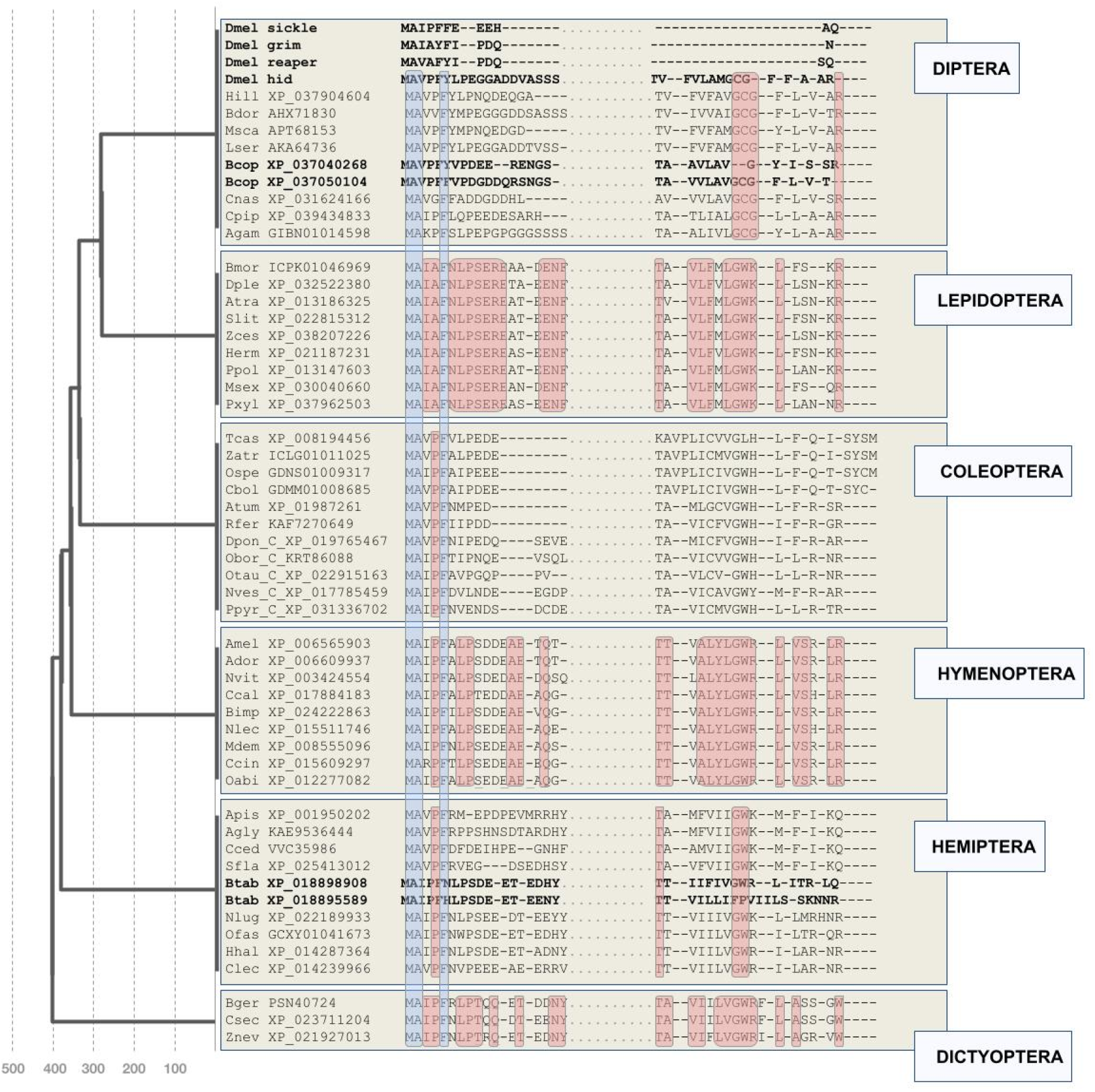
Overview on newly detected RHG homologs. Clustal Omega multiple sequence alignment oF N- and C-terminal regions for a selection of newly RHG identified homologs. Blue overcasts: Residues conserved across all homologs. Red overcasts: Residues conserved across all homologs within orders (except within-species paralogs). Duplicated homologs in *D. melanogaster* (*sickle, grim, reaper, hid*), the fungus gnat *Bradysia odoriphaga* (Brad), and *Bemisia tabaci* (Btab) indicated by bold font. See Supplementary data file 1 for species abbreviations. Numbers at the bottom of hatched vertical time lines correspond to millions of years past present time. Divergence time points based on Misof et al. (2014).

Most of the newly detected homologs outside the genus *Drosophila* were singletons except for duplicate pairs in the silverleaf whitefly *Bemisia tabaci* and the fungus gnat *Bradysia odoriphaga* (Fig. 1), and the exceptional expansion of RHG homologs in mosquitoes (see below).

### Protein sequence conservation differences within and between orders

The crucial success of “taxon-hopping” in the detection of new RHG homologs constituted preliminary evidence of potentially different rates of RHG sequence change between and within insect orders. This idea was further supported by the clade-specific differences of sequence divergence in the most conserved protein sequence regions of the newly detected RHG homologs, i.e. the N- and C-terminal ends (Fig. 1). To test for this possibility in a quantitative manner, I generated estimates of RHG protein sequence change rates within insect orders by determining average numbers of non-conserved sites in Clustal Omega multiple sequence alignments (MSAs) divided by respective clade ages (Table 1). By this measure, RHG protein sequence change rates varied up to over 15-fold between select clades. The lowest rate was found for the hymenopteran RHG homologs with 0.18% per million years while aphids stood out with the highest rate of close to 3% per million years (Table 1). These outliers excluded, the average RHG protein sequence change rate amounted to 0.34% (+/−0.07) per million years. Further notable was the fact that the aphid protein sequence change rate of 3% per million years compared to 0.29% in the remaining hemipteran species sampled (Table 1). Thus, while approximate, the quantitative findings confirmed the existence of RHG protein sequence change rate differences between and within insect orders.

**Table 1:** Clade specific diversification rates of RHG homologs. The dipteran sample included *D. melanogaster hid* but no homologs of *skl*, *grim*, or *rpr*. See Supplementary data files 3-9 for corresponding MSAs. Divergence times based on Misof et al. (2014).

|  | Average<br>lengths | % Conserved sites | Divergence<br>times | % divergence/million years |
| --- | --- | --- | --- | --- |
| **Diptera (n=13)** | 302.0 | 17.2% | 200 | 0.41% |
| **Lepidoptera (n=13)** | 210.1 | 54.7% | 120 | 0.38% |
| **Coleoptera: Polyphaga (n=24)** | 189.3 | 4.2% | 250 | 0.38% |
| **Hymenoptera (n=19)** | 244.9 | 54.3% | 250 | 0.18% |
| **Hemiptera wo Aphidoidea (n=8)** | 225.5 | 24.4% | 260 | 0.29% |
| **Aphidoidea (n=10)** | 252.8 | 25.3% | 25 | 2.99% |
| **Dictyoptera (n=3)** | 253.3 | 56.1% | 175 | 0.25% |

### Deeply conserved N- and C-terminal end amino acid residues

Despite the partly dramatic differences in protein sequence divergence, MSAs of the newly compiled RHG protein sequences also identified deeply conserved amino acid residues. This was not only the case for the previously noted conserved N-terminal IBM, but also for residues at the C-terminal end (Fig. 1). Most consistent was the deployment of arginine (R) as the terminal amino acid residue, which is also the case for *Drosophila* RHG homolog *hid* (Fig. 1). Besides the Drosophila RHG paralogs *skl*, *grim*, and *rpr*, other exceptions included the duplicated RHG homologs of *B. tabaci* and *B. odoriphaga*. Moreover, in all of the compiled aphid homologs, the ancestral arginine residue was replaced by glutamine (Q), a feature shared by one of the duplicated homologs in the closely related *B. tabaci* (Fig. 1).

A second example of clade-specific departure from the N-terminal amino acid residue consensus were the exceptionally sequence diverged N-termini in a subgroup of darkling beetles including *T. castaneum* (Fig. 1). The three dictyopteran species, finally, shared a terminal tryptophan residue (Fig. 1).

The second-most consistently conserved C-terminal pattern was the combination of a glycine (G) residue followed by tryptophan (W) 5-7 residues away from the C-terminus in all clades except Diptera (Fig. 1). The latter shared the conserved glycine residue but the adjacent consensus tryptophan was replaced by a cysteine (C). Further exceptions from the GW consensus included the RHG homolog of *T. castaneum* and one of the two *B. tabaci* paralogs, XP_018895589, which lacked both residues (Fig. 1).

There was also tentative evidence of protein sequence conservation further N-terminal of the conserved glycine-tryptophan duplet, which was more unambiguously documented in the sequence comparisons within orders than between orders (Fig. 1). Overall, however, these findings unearthed evidence of deeply conserved constraints at the C-terminal end of RHG proteins in addition to the N-terminal IBM. Moreover, these findings also defined *hid* as the most ancestrally organized RHG paralog in *Drosophila* given the complete lack of C-terminal consensus residues in *rpr*, *grim*, and *skl* (Fig. 1).

### *michelob_x* constitutes an independent RHG gene family expansion in mosquitoes

The first RHG homologs outside the genus *Drosophila* were discovered in mosquitoes (Zhou et al. 2005). Completion of the genome sequence project of *A. gambiae* revealed the presence of conserved caspase and IAP genes but the existence of RHG homologs had initially remained elusive (Christophides et al. 2002; Zdobnov et al. 2002). Developing a hidden Markov model (HMM) search profile for the RHG IBM motif from sequence comparisons of distantly related *Drosophila* species, Zhou et al. (2005) detected candidate RHG homologs in *A. gambiae*. One of these, named *michelob_x* (*mx*), was studied in detail and found to induce apoptosis in cell culture as well as transgenic *Drosophila*. Moreover, *A. gambiae* Mx was also shown to bind Drosophila Diap1 in vitro and in an IBM dependent manner (Zhou et al. 2005).

Given the apparent lack of *hid*-like C-terminal consensus amino acid residues in mosquito *mx* homologs (Fig. 2) (Zhou et al. 2005), I conducted BLAST searches against mosquito genome and transcript databases with both *mx* and dipteran *hid*-like RHG homologs as queries. These efforts revealed the presence of *hid*-like RHG homologs in *A. gambiae* and other mosquito species (Fig. 1 and Supplementary data file 1). Moreover, while no *mx*-like homologs were detectable outside the dipteran suborder Culicomorpha, two additional *mx*-like paralogs were found in members of the mosquito subfamilies Culicinae (*Aedes aegypti*, *Culex pipiens*, *Tripteroides aranoides*) and Toxorhynchitinae (*Toxorhynchites* spec.) (Fig. 2). Combined, these findings uncovered an expansion of the derived *mx*-type RHG subfamily in mosquitoes, paralleling that of *rpr*, *grim*, and *skl* in the higher Diptera. Unlike in the latter case, however, the C-termini of the mosquito *mx* paralogs were characterized by a high degree of overall sequence conservation with tyrosine (Y) as the C-terminal residue (Fig. 2 and Supplementary data file 2).

**Figure 2:**
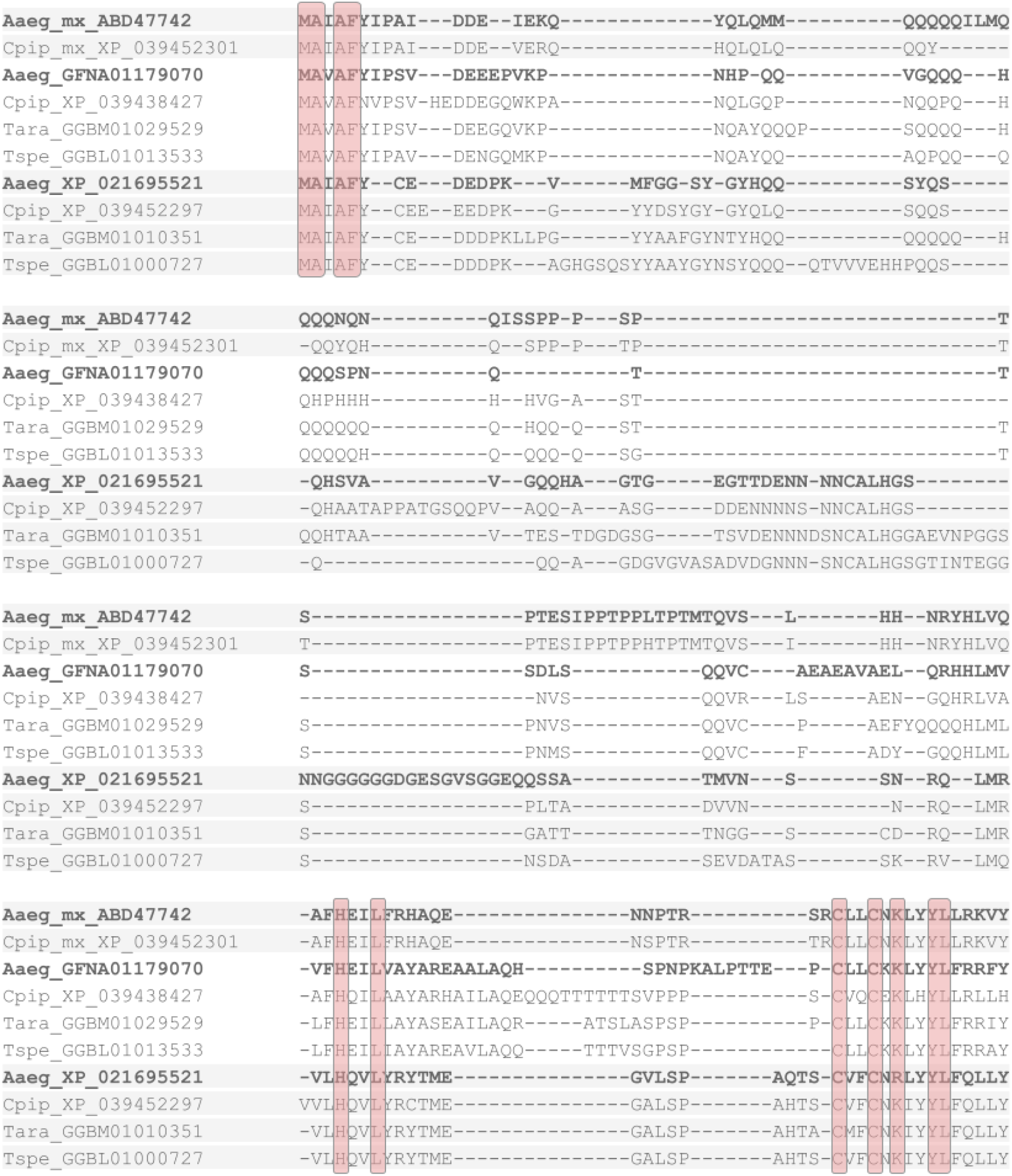
Multiple sequence alignment of mosquito *mx* paralogs. Multiple sequence alignment of *mx* homologs detected in *Aedes aegypti* (Aaeg), *Culex pipiens* (Cpip), *Tripteroides aranoides* (Tara), and *Toxorhynchites* spec. (Tspe). Residues conserved across all homologs highlighted by red overcast. *A. aegypti* homologs are highlighted in bold font for orientation.

### Gene structure conservation

The open reading frames (ORFs) of *Drosophila grim*, *rpr*, and *skl* are localized on single exons, while the ORF of *hid* spreads out over 4 exons (Grether et al. 1995), an organization that is conserved in the *hid* ortholog of the scuttle fly *Megaselia scalaris* (Yoo et al. 2017). To probe for possibly conserved gene structures in the newly identified RHG homologs, I investigated exon-intron organization of 15 newly identified homologs based on transcript expression (RNAseq) supported gene models in the gene database of NCBI (Fig. 3). RHG homolog selection was guided by covering maximal phylogenetic depth for each order and included experimental model systems such as the silkworm moth *B. mori* (Meng et al. 2017), the red flour beetle *T. castaneum* (Brown et al. 2009), the jewel wasp *Nasonia vitripennis* (Werren and Loehlin 2009), and the milkweed bug *Oncopeltus fasciatus* (Chipman 2017).

**Figure 3:**
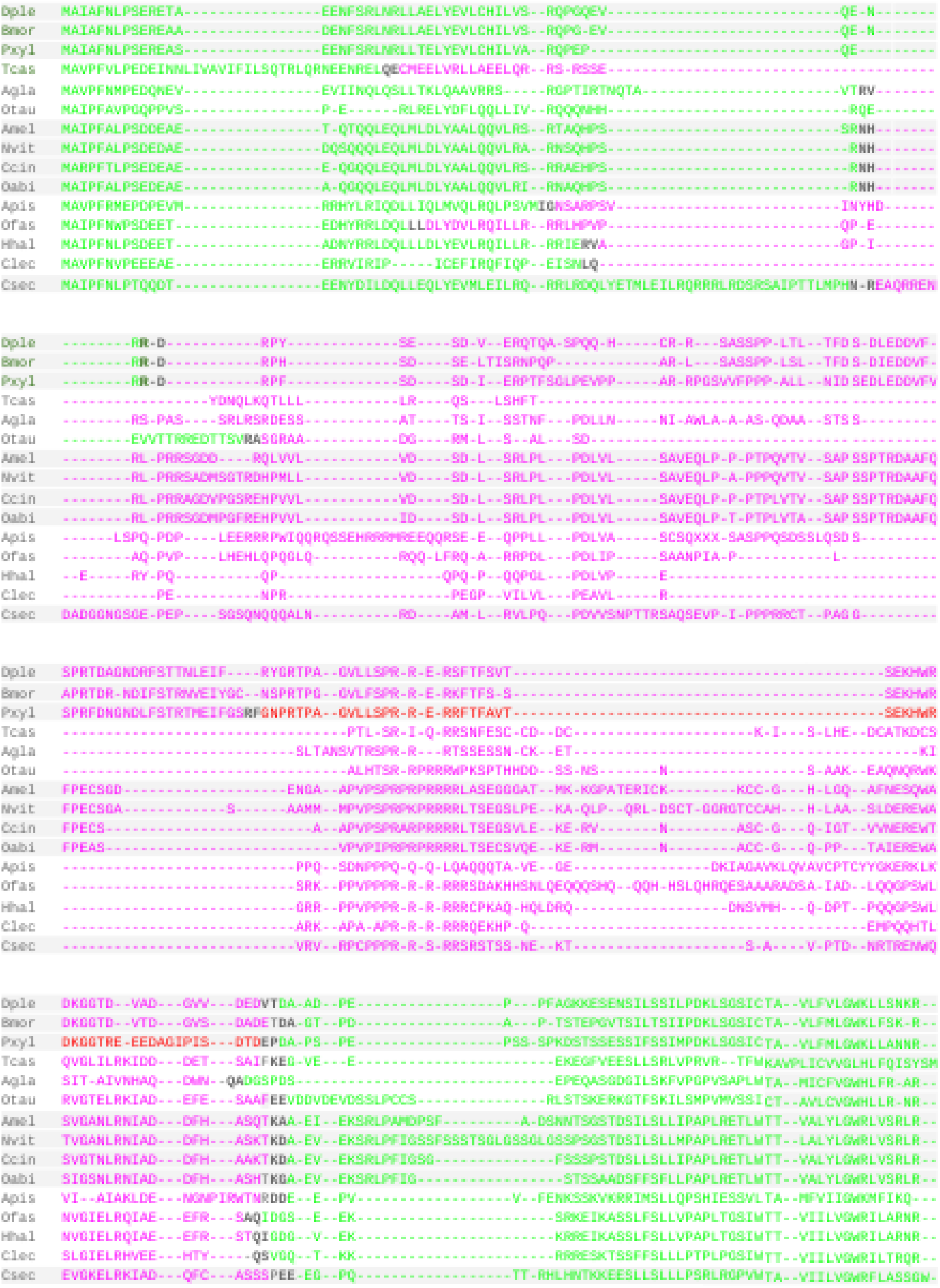
Gene structure conservation. Multiple sequence alignment of representative, newly identified RHG homologs. Exon borders indicated by black bold font. Sequence from different exons sequentially colorized green and purple. Additional exon in *P. xylostella* is highlighted in red. Light grey background shades indicate different insect orders.

In the great majority of cases, ORFs were spread out over three exons (Fig. 3). Splicing site positions, however, were only conserved in a few cases within orders, most obviously in the Lepidoptera and Hymenoptera. In the former, the homolog of the oldest clade sampled, i.e. the Yponomeutoidea represented by the diamondback moth *Plutella xylostella*, was characterized by the acquisition of an exceptional fourth ORF encoding exon (Fig. 3).

In general, the N- and C-terminal ORF segments were encoded on smaller exon contributions than the intermittent regions, which also differed by a higher level of sequence divergence. Moreover, the ORF position of the splice site linking the intermittent region with the C-terminus appeared generally more strongly conserved than the positions of other splice sites. Thus, overall, the RHG homologs were characterized by a deeply conserved gene organization resembling that of Drosophila *hid* (Grether et al. 1995).

### A conserved RHG isoform in the Lepidoptera

In many Lepidoptera, initial BLASTp searches recovered two types of RHG orthologs per species. In these cases, the two apparent homologs were sequence identical in the N-terminal region but diverged C-terminally. This preliminary evidence of differential splice isoforms was confirmed by the gene structure analyses. In the genome draft of the Monarch butterfly, *Danaus plexippus*, for instance, one isoform (OWR53643.1) was identified among the curated protein sequence predictions (Zhan et al. 2011) and a second (XP_032522380) among the automatic protein sequence predictions in the genome daft assembly Dplex_v4 (GenBank assembly accession: GCA_009731565.1). The same organization was eventually found for all sampled lepidopteran homologs. One isoform was the product of run-off translation from the first exon. As the resulting predicted proteins were on average 80 amino acids shorter than those of the second isoform resulting from the 2-3 exons spanning ORFs, it seemed appropriate to name the two isoforms short (S) and long (L) RHG isoforms, respectively (Fig. 4). The fact that both isoforms were also found in the diamondback moth *P. xylostella*, i.e. the representative of the Yponomeutoidea, implies at least 140 million years of evolutionary conservation of both isoforms in the Lepidoptera (Kawahara et al. 2019).

**Figure 4:**
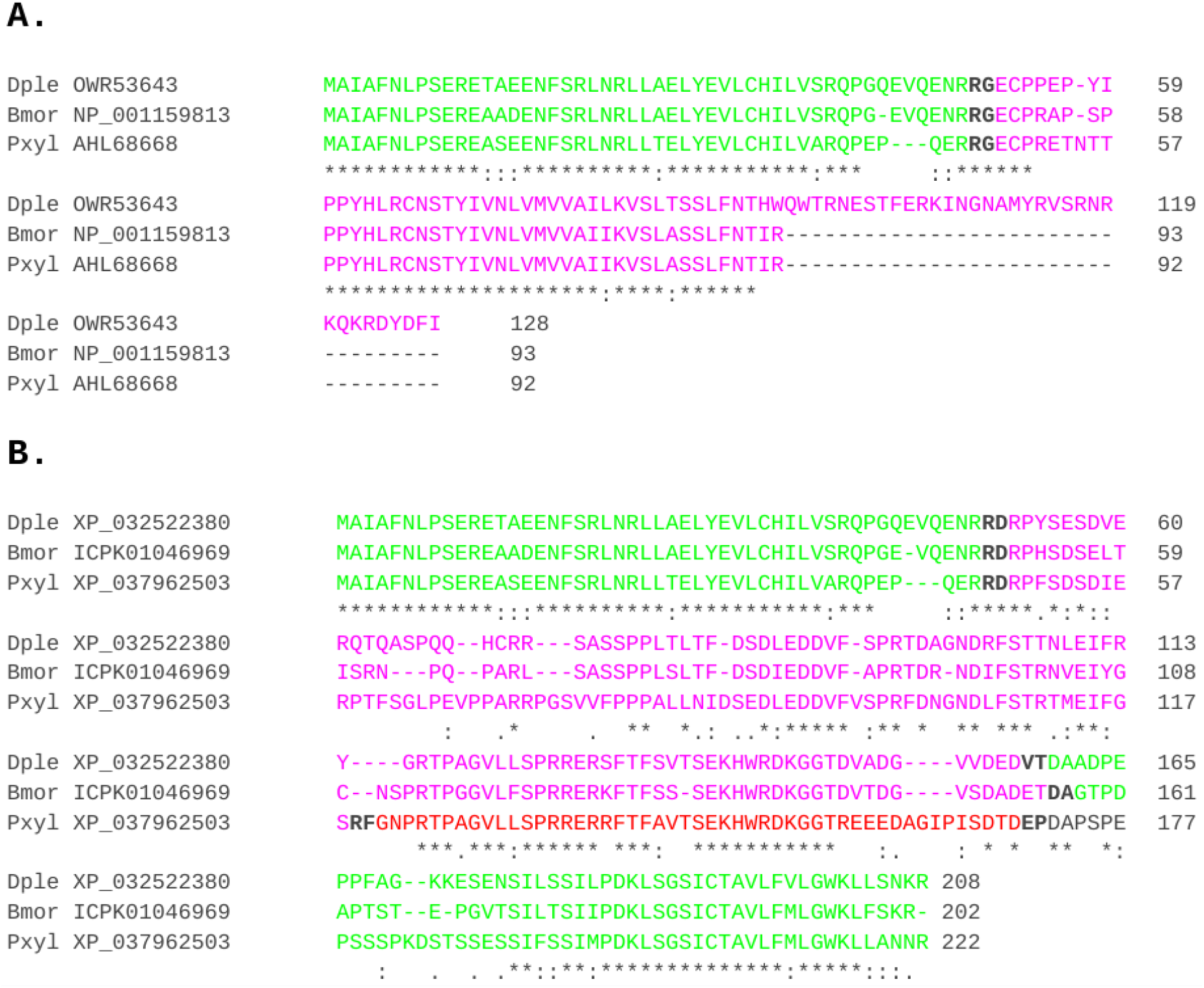
RHG protein products of conserved splice isoforms in the Lepidoptera. **(a)** Multiple sequence alignment of the RHG S-isoforms of *D. plexippus* (Dple), *B. mori* (Bmor), and *P. xylostella* (Pxyl). (**a)** Multiple sequence alignment of the L-isoform protein sequences for the same species. Exon boundary highlighting and protein sequence color coding in as in Fig. 2.

### Exceptional RHG sequence divergence in aphids

Past efforts failed to identify RHG homologs in the pea aphid *A. pisum*, an important pest species and genome evolution model (International Aphid Genomics Consortium 2010; Ribeiro Lopes et al. 2019; Julca et al. 2020). Using the C-terminal RHG homolog region of the brown planthopper *Nilaparvata lugens* as query in a PSI-BLAST search against the nr database for the taxonomic range of aphid species (Aphidoidea) yielded a single hit in the yellow sugarcane aphid, *Sipha flava*, with an e value of 0.009. Subsequent searches with the *S. flava* RHG homolog uncovered single copy hits in nine additional aphid species including *A. pisum* (Fig. 1, 5a, and Supplementary data file 1). Most of the aphid homologs were characterized by a number of glutamine (Q) and proline (P) repeat strings in the middle region of the protein, some of which were of variable lengths even between closely related species. Similar repetitive sequence elements were also found in other hemipteran RHG homologs (Fig. 5a). The protein sequence of *A. pisum*, however, stood out by a unique 13 repeat units long string of the sextamer “(S/H)(A/V)GP(S/L/P)(H/Q)” with six perfect copies of “SAGPSH” (Fig. 5a and b). Expression of this simple sequence region was supported by RNAseq data mapped against the gene *A. pisum* RHG gene model in the NCBI gene database (not shown). Similarity blotting of the *A. pisum* RHG coding sequence confirmed corresponding repetitiveness as the nucleotide level, which is typical for slippage extended simple sequence repeats (Fig. 5b) (Schlötterer and Tautz 1992).

**Figure 5:**
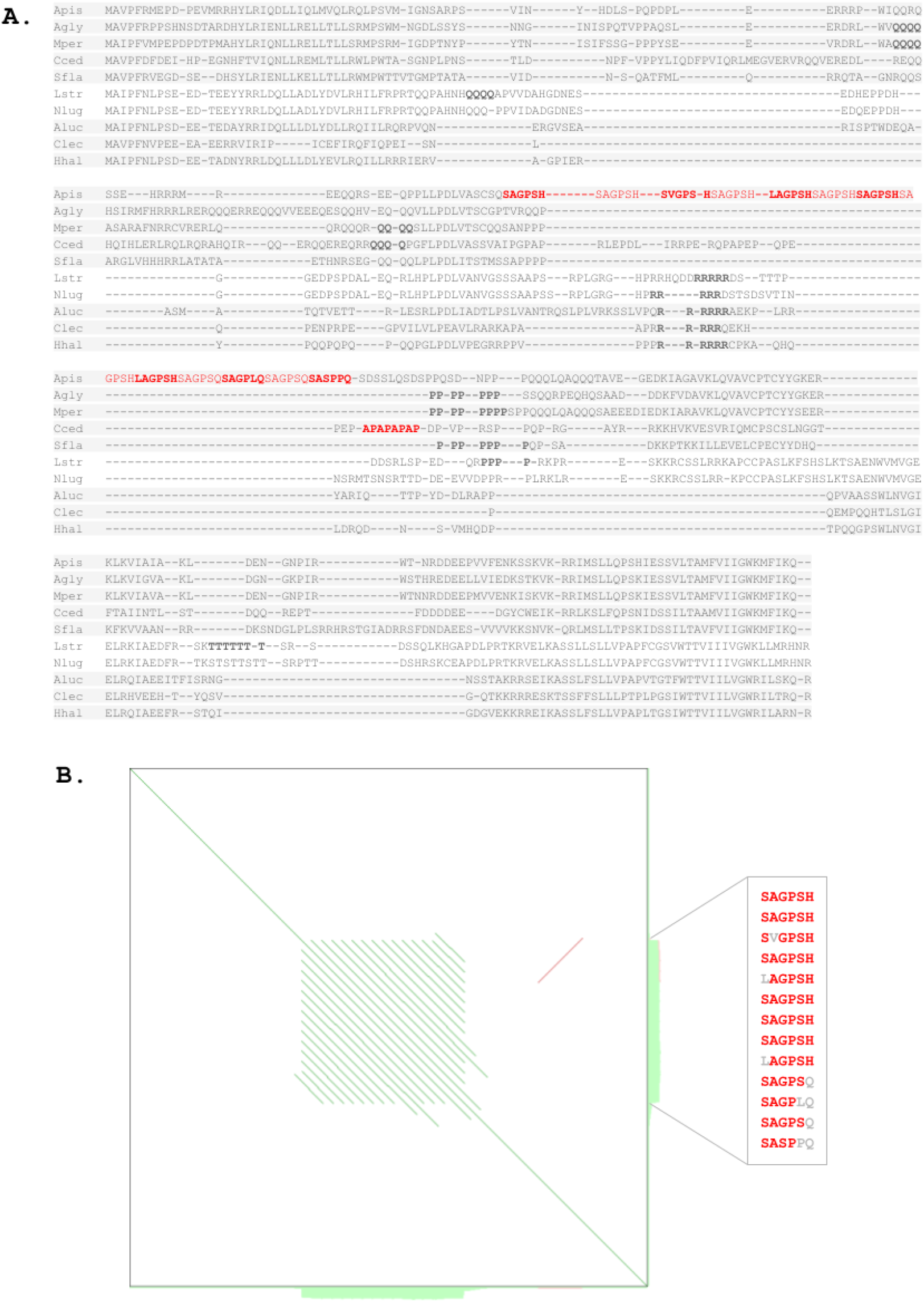
Protein sequence divergence in aphid RHG homologs. (**a)** Multiple sequence alignment of hemipteran RHG homolog protein sequences. Background shade visualizes clade composition. Top 5 species represent members of the family Aphididae (Apis = *Aphis pisum*, Agly = *Aphis glycines*, Mper = *Myzus persicae*, Cced = *Cinara cedri*, Sfla = *Sipha flava*). Species number 6 and 7 from the top are planthoppers (Auchenorrhyncha: Lstr = *Laodelphax striatellus*, Nlug = *Nilaparvata lugens*). Bottom 3 species represent the suborder Heteroptera (Aluc = *Apolygus lucorum*, Clec = *Cimex lectularius*, Hhal = *Halyomorpha halys*). Single amino acid repeat strings longer than 3 residues highlighted in bold font. 13-mer repeat of the hexapeptide “(S/H)(A/V)GP(S/L/P)(H/Q)” in the pea aphid and strong of the residue duplet AP in *Cinara cedri* highlighted in red font. **(b)** Sequence similarity dot blot generated with YASS (Noé and Kucherov 2005) of the *A. pisum* RHG coding region DNA sequence XM_001950167.5 visualizing internal repetitiveness of the 13-mer “(S/H)(A/V)GP(S/L/P)(H/Q)” repeat at the nucleotide sequence level. Green shading along blot edges indicate significantly repetitive sequence regions. Box to the right shows alignment of the 13 repeats stacked top to bottom in N- to C-terminal direction with main consensus residues highlighted by bold red font and variant residues indicated by grey font.

A second unusual characteristic of the aphid RHG homologs was their consistent deployment of glutamine (Q) as the N-terminal residue in place of the deeply conserved ancestral arginine (R) residue in the Hemiptera and other insect orders (Fig. 1 and 5a). Combined, the stronger departure of aphid RHG homologs from some of the broadly conserved RHG sequence characteristics provided an explanation for their lower detectability with query sequences from distantly related species.

## Discussion

The expanded panel of insect RHG homologs clarifies a number of previously elusive aspects of this critical cell death gene family. Most importantly, perhaps, and consistent with previous speculations (Yoo et al. 2017), *hid* is now clearly established as the most ancestrally organized member of the four *Drosophila* RHG paralogs via outgroup comparison. Further significant, the protein product of *hid*, in contrast to *rpr*, *skl*, and *grim*, is localized to mitochondria due to its hydrophobic C-terminus (392-409), which has therefore been defined as the mitochondria-targeting sequence (MTS) domain (Haining et al. 1999). Thus, while the role of mitochondria in *Drosophila* cell death is still not clearly defined, the conservation of N-terminal residues, i.e. a *hid*-like MTS domain, in ancestral RHG homologs across winged insects further suggests that mitochondrial localization is a critical aspect of insect RHG protein function. The possibility that the MTS domain of *hid* promotes IAP degradation by virtue of mitochondrial colocalization therefore continues to be an attractive model (Sandu et al. 2010; Yoo et al. 2017). This is further supported by the fact that both the IMB and MTS domain are essential for Hid’s cell death inducing capacity (Yoo et al. 2017). Interestingly, also the lepidopteran S-isoform is mitochondrially localized based on immunohistochemical detection in the armyworm moth *S. frugiperda* (Shu et al. 2020), suggesting at a higher level of functional conservation between the derived S- and ancestrally organized L-isoforms in the Lepidoptera compared to that between *hid* vs *grim*, *rpr*, and *skl* in *Drosophila*.

The updated insect RHG homolog database further reveals that *rpr*, *skl*, and *grim* are not the only examples of RHG gene family expansions resulting into paralogs with simpler gene organization, i.e. a lower number of coding exons, and substantially shorter protein sequences. This is also true for the *mx* paralogs in mosquitoes and the derived S-isoforms of the lepidopteran RHG genes. The discovery of the latter further suggests that the dipteran RHG gene family expansions occurred via selective duplication of the first ORF sequence containing exon of the ancestrally organized singleton precursor RHG encoding the short, but cell death induction sufficient, IBM domain. This duplication conduciveness explains the spawning of short, compact paralogs like *rpr*, *grim*, and skl or the *mx* paralogs in mosquitoes from ancestral hid-like RHG precursor genes (Yoo et al. 2017).

Interestingly, the existence of multiple *mx* homologs had been noted earlier (Wang and Clem 2011). Tissue- and, ideally, cell-specific expression studies will reveal whether and how these duplications translated into functional diversification of the mosquito *mx* subfamily in relation to the ancestrally *hid*-like homologs in mosquitoes, which await to be studied. While these efforts may reveal connections to the exceptional pathogen load of mosquito vector pest species, it is also possible that they represent functionally neutral outcomes of gene duplication in line with the “duplication-degeneration-complementation” trajectory (Force et al. 1999; Oakley et al. 2006). This, in fact, could apply to *hid*, *rpr*, *grim* and *skl*, given their largely overlapping expression dynamics based on modENCODE data (Chen et al. 2014).

The first RHG homologs identified outside dipterans via bioinformatic search in a new genome sequence was *Ibm1* of *B. mori* (Bryant et al. 2009), which is now identified as the derived S-isoform of the *B. mori* RHG homolog locus. Paralleling the situation in mosquitoes, it is the ancestrally organized L-RHG isoform that has to be studied (Bryant et al. 2009; Shu et al. 2020). Future analyses of both lepidopteran isoforms have the potential to inform about the subfunctionalization trajectories of newly emerging RHG homologs. In this case, the existence of post-transcriptional mechanisms can be envisioned to confer cell- or tissue-specific functions.

It has been over 10 years since the last RHG homolog was detected in a new insect order. This hiatus is in part explained by the well recognized challenges of finding RHG homologs, i.e. their short sequence lengths, relatively unconstrained evolution, and low number of constrained residues. However, the updated RHG compilation also reveals a role of historical contingencies. The previously identified homologs in mosquitoes and lepidopterans both represent derived homologs or isforms, lacking the conserved C-terminus. This may, in part, explain the failures of previous efforts such as in Hemipterans, for example (Ribeiro Lopes et al. 2019). The latter case, however, is also an example of yet another likely impeding coincidence. Some ancestrally organized RHG homologs have exceptionally diverged even in the N- and C-terminal regions, thus reducing their detectability. This is true for aphids, including *A. pisum*, arguably the genomically best documented representative of its clade (International Aphid Genomics Consortium 2010). The second example of an advanced genomic model organism possessing an exceptionally diverged RHG homolog is the red flour beetle *T. castaneum*. The RHG homologs of both species were only detected after homolog sequences from within the same insect order were at hand as queries, i.e. through “taxon hopping”.

Varied BLAST searches in genome and transcriptome databases of older insect clades, i.e. Paleoptera and Apterygota, as well as crustaceans and invertebrates in general did not uncover further RHG homologs at this point. Given the success of the “taxon hopping” strategy in identifying new homologs, it seems reasonable to assume that the RHG gene family is restricted to insects, if not specifically winged insects. Thus, besides identifying new powerful insect model systems for the study of RHG function, the expanded compilation of RHG homologs provides a new hypothetical time point of RHG family origination at the base of pterygote insects, predicting the existence of different IAP inhibiting regulators in other clades.

## MATERIALS AND METHODS

### Homolog searches

Using the BLAST search interface of the National Center for Biotechnology Information (NCBI), Homolog searches were conducted with BLASTp, tBLASTn or Position-Specific Iterated BLAST (PSI-BLAST) in the non-redundant (nr) protein sequence, Transcriptome Shotgun Assembly (TSA), and Whole genome shotgun contig (wgs) sequence databases (Altschul and Koonin 1998; McGinnis and Madden 2004; Pruitt et al. 2005). Most searches were performed at default settings. In rare cases, searches were repeated with setting word size to 3 and expected threshold to 0.5.

### Multiple sequence alignments

Multiple protein sequence alignments were generated using Clustal Omega, webPRANK, and T-Coffee all at default settings (Notredame et al. 2000; Löytynoja and Goldman 2010; Sievers et al. 2011).

### Gene structure analyses

Gene structures were analyzed in current assemblies made available by the NCBI Assembly database (Pruitt et al. 2005).

## ACKNOWLEDGEMENTS

I thank Lori Pile and the anonymous reviewers for helpful comments on the manuscript.

**Supplementary data file 1:** Protein sequence IDs and species information of RHG homolog compilation.

**Supplementary data file 2:** MSA of culicomorph RHG protein sequences. Top five sequences: *mx* homologs from all investigated mosquito species. Red and green font: Additional *mx* homologs in species from the subfamily Culicinae. Hid-like homologs from mosquito and other dipteran species. See Supplementary data file 1 for species abbreviations.

**Supplementary data file 3**: MSA of dipteran RHG protein sequences. See Supplementary data file 1 for species abbreviations.

**Supplementary data file 4:** MSA of lepidopteran RHG protein sequences. See Supplementary data file 1 for species abbreviations.

**Supplementary data file 5:** MSA of coleopteran RHG protein sequences. See Supplementary data file 1 for species abbreviations.

**Supplementary data file 6:** MSA of hymenopteran RHG protein sequences. See Supplementary data file 1 for species abbreviations.

**Supplementary data file 7:** MSA of hemipteran RHG protein sequences, Aphidoidea excluded. See Supplementary data file 1 for species abbreviations.

**Supplementary data file 8:** MSA of Aphidoidea RHG protein sequences. See Supplementary data file 1 for species abbreviations.

**Supplementary data file 9:** MSA of dictyopteran RHG protein sequences. See Supplementary data file 1 for species abbreviations.

Supplementary data file 10: Amino acid sequences of all homologs analyzed in FASTA format.

## Notes

https://drive.google.com/drive/folders/1uR5UkAonZePkePO3TUdGGwVj3WMuauOC?usp=sharing

### Competing Interest Statement

The authors have declared no competing interest.

